# Novel transformer networks for improved sequence labeling in genomics

**DOI:** 10.1101/836163

**Authors:** Jim Clauwaert, Willem Waegeman

## Abstract

In genomics, a wide range of machine learning methodologies have been investigated to annotate biological sequences for positions of interest such as transcription start sites, translation initiation sites, methylation sites, splice sites and promoter start sites. In recent years, this area has been dominated by convolutional neural networks, which typically outperform previously-designed methods as a result of automated scanning for influential sequence motifs. However, those architectures do not allow for the efficient processing of the full genomic sequence. As an improvement, we introduce transformer architectures for whole genome sequence labeling tasks. We show that these architectures, recently introduced for natural language processing, are better suited for processing and annotating long DNA sequences. We apply existing networks and introduce an optimized method for the calculation of attention from input nucleotides. To demonstrate this, we evaluate our architecture on several sequence labeling tasks, and find it to achieve state-of-the-art performances when comparing it to specialized models for the annotation of transcription start sites, translation initiation sites and 4mC methylation in *E. coli*.

## 1 Introduction

In the last 30 years, a major effort has been invested into uncovering the relation between the genome and the biological processes it interacts with. A thorough understanding of the influence of the DNA sequence is of importance for the manipulation of biological systems, e.g. to facilitate the forward engineering of biological pathways. In recent years, machine learning methodologies play an increasingly import role in the construction of predictive tools. These tasks include the annotation of genomic positions of relevance, such as transcription start sites, translation initiation sites, methylation sites, splice sites and promoter start sites.

Early methods for labeling of the DNA sequence were focused on the extraction of important features to train supervised learning models, such as tree-based methods or kernel methods. More recently, convolutional neural networks (CNNs) have been popular, initiated from the work of Alipanahi et al. [1]. The popularity of the CNN can be attributed to the automatic optimization of motifs or other features of interest during the training phase.

The prokaryotic and eukaryotic genome is built from 10^7^ and 10^10^ nucleotides. Given its size, only a small fragment of the genome sequence is bound to determine the existence of certain genomic sites. In order to create a feasible sample input, only a short fragment of the genome sequence is used to predict the occurrence of these sites. The boundaries of this region with respect to the position of interest is denoted as the receptive field. Due to the model architecture of conventional machine learning and deep learning techniques such as convolutional neural networks, where the input is structured according to the relative distances towards the output label, custom input samples are created from the genome, in accordance to the selected receptive field around each nucleotide position. However, input samples of neighboring positions are created from largely overlapping regions on the genome. When evaluating all positions on the genome, the combined sequence length of the input samples is several times larger than the length of the original genome, and scales with the size of the receptive field.

In practice, existing studies do not apply the full genome for training or evaluation. This task is too resource-heavy for a multitude of machine learning methodologies that have not been created to handle millions of samples. Additionally, the majority of the annotation tasks represent a positive and negative set that is heavily imbalanced, which can hinder the success of learning approaches. For example, the detection of transcription start sites (TSSs) has several thousand times more negative than positive labels. In some cases, the site of interest is constrained to a subset of positions. This is exemplified by the site at which translation of the RNA is initiated, denoted as the Translation Initiation Site (TISs), where valid positions can be delimited by three nucleotides being either ATG, TTG or GTG [7]. For annotation tasks that can not be constrained to a smaller set, the negative set is sampled (e.g. prediction of TSS [23] [34] or methylation [17]). In general, the size of the sampled negative set is chosen to be of the same order of magnitude as the size of the positive set, constituting only a fraction of the original negative set size (0.01% for TSS in *E. coli*). However, given the comparative sizes of the sampled and full negative set, performance metrics are not guaranteed to correctly reflect the models predictive capability. When considering the task for which the model is optimized, it is plausible that the resulting performances generalize poorly when applying the model on the full genome.

Transformer networks have recently been introduced in natural language processing [35]. These architectures are based on attention and outperform recurrent neural networks on natural language sequence-to-sequence labeling benchmarks. In 2019, Dai et al. [9] defined the transformer-XL, an extension of the transformer unit for tasks constituting long sequences through introduction of a recurrence mechanism, showing promise towards evaluating the genome sequence. In this study, we introduce a novel transformer-based model for DNA sequence labeling tasks. The genome is processed as is, where nucleotide inputs contribute to the prediction of multiple outputs. The size of the receptive field does not influence the amount of data processed, nor does it influence the amount of parameters of the model. In contrast to recurrent neural networks, which share these advantageous properties, the transformer-XL architecture iterates the genome sequence in segments of multiple nucleotides, offering superior processing times. By applying a model on the full genome, no complications arise that are linked to subsampling of the negative set.

We define for the first time a transformer architecture for DNA sequence labeling by building upon recent innovations in the field of natural language processing. Second, we substantiate and implement adaptations to the model that make it better suited to extract information from nucleotide sequences, an extension that proves to drastically improve the predictive capabilities of the model. Third, a benchmark is performed with recent studies for three different annotation tasks: transcription start sites, translation initiation sites and methylation sites. We prove that the novel transformer network attains state-of-the-art performances, while retaining fast training times.

## 2 Related work

Studies exploring data methods for statistical inference based solely on the nucleotide sequence go back as far as 1983, with Harr et al. [13] publishing mathematical formulas on the creation of a consensus sequence for TSSs in *E. coli*. Stormo [32] describes over fifteen mathematical approaches in relation to processing DNA sequences between 1983 and 2000, ranging from: algorithms designed to identify consensus sequences [30] [21], tune weight matrices [33] and rank alignments [19, 41].

With the increased knowledge in the field of molecular biology and the failing attempts to create robust correlations between the DNA sequence and properties of interest, efforts towards feature engineering were made, extracting physical, chemical and biological meaning that could show relatedness towards the biological process. Several important descriptors of sequences include, but are not limited to: the GC-content, bendability [42], flexibility [3] and free energy [16]. Recently, Nikam et al. published Seq2Feature, an online tool that can extract up to 252 protein and 41 DNA sequence-based descriptors [27].

The rise of novel machine learning methodologies, such as Random Forests and support vector machines, have resulted in many applications for the creation of tools to annotate the prokaryotic genome. Liu et al. propose stacked networks that apply Random Forests [24] for two-step sigma factor prediction in *E. coli*. Support vector machines are applied by Manavalan et al. to predict phage virion proteins present in the bacterial genome [25]. Further examples of the application of support vector machines include the work of: Goel et al. [12], who propose an improved method for splice site prediction in Eukaryotes; and, Wang et al. [36], who introduce the detection of *σ*^70^ promoters using evolutionary driven image creation.

Successful gains in the field of machine learning and genome annotation can be attributed to the use of deep learning methods. In 1992, Horton et al. [14] published the use of the first perceptron neural network, applied for promoter site prediction in a sequence library originating from *E. coli*. However, the popular application of deep learning started with CNNs, initially designed for networks specializing in image recognition. These incorporate the optimization (extraction) of relevant features from the nucleotide sequence during the training phase of the model. Automatic training of position weight matrices has achieved state-of-the-art results for the prediction of regions with high DNA-protein affinity [1] in eukaryotes. As of today, several studies have been applying CNNs for prokaryotes. These include models for the annotation of methylation sites [11], origin of replication sites [10] [22], recombination spots [5][39], TSSs [34].

Recurrent neural network architectures have not been applied on the full genome due to their long processing times, but can be used to process individual samples of decreased lengths featuring expression and/or nucleotide sequence data. In combination with convolutional layers, they have been to used to detect TISs [7] for *E. coli*. More applications of recurrent neural networks exist for Eukaryotes, such as miRNA target prediction [20] and, combined with convolutional layers, the annotation of methylation states [37, 2] and detection of protein binding sites on RNA [15].

The only type of machine learning method that has been successfully applied on the full genomic sequence are hidden Markov models. However, due to the limited capacity of a hidden Markov model, this method is nowadays rarely used for new studies. Some applications for prokaryotes include the detection of genes in *E. coli* [18] and the recognition of repetitive DNA sequences [38].

## 3 Transformer Network

Here we describe our transformer network for DNA sequence labeling. In Section 3.1, we adapt the transformer architecture of Dai et al. [9] to DNA sequences. Unlike the natural language processing tasks for which the transformer-XL architecture was first described, the annotation of DNA sequences is not an autoregressive task. Additionally, only a few input and output classes exist, resulting in the models being less complex. To better extract features from the nucleotide sequence, an adaptation to the calculation of attention is described in Section 3.2.

In this paper, we adopt the annotation of normal lowercase letters for parameters (e.g. *l*), bold lowercase letters for vectors (e.g. **q**) and uppercase letters for matrices (e.g. *H*).

### 3.1 Basic model

In essence, the annotation of DNA is a sequence labeling task that has correspondences to natural language processing. Representing a DNA sequence of length *p* as (*x*_1_, *x*_2_,…, *x_p_*), where *x_i_* ∈ {*A, C, T, G*}, the tasks described in this paper are binary classification problems, with each position *x_i_* having a corresponding label *y_i_* ∈ {0, 1}. A positive label denotes the occurrence of an event at that position.

The model processes the genome in sequential segments of *l* nucleotides. During training, a nonlinear transformation function *E* is optimized that maps the input classes {*A, C, T, G*} to a vector embedding **h** of length *d_model_*. for nucleotide *x_i_* on the genome:

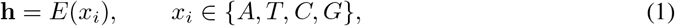

where 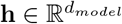.

The inputs at each segment are processed through *k* layers. Within each layer, multi-head attention is calculated for each hidden state **h** using the collection of hidden states [**h**^(0)^…**h**^(*l*−1)^] within each segment, represented as rows in the matrix 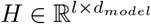.

Next, for each hidden state of **h**, the output of the multi-head attention step (*MultiHead*) is summed with the input, i.e. a residual connection. The final mathematical step within each layer is layer normalization [4]. The operations for hidden states **h** in layer *t* at position *n* in segment *s* are performed in parallel:

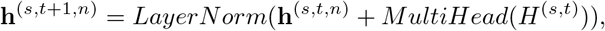

or,

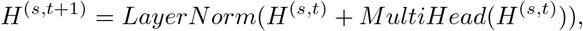

where *t* ∈ [0, *k*[and *n* ∈ [0, *l*[.

After a forward pass through *k* layers, a final linear combination reduces the dimension of the output hidden state (*d_model_*) to the amount of output classes. In this study, only binary classification is performed. A softmax layer is applied before obtaining the prediction value 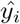 for nucleotide *x_i_*.

#### 3.1.1 Multi-head attention

The core functionality of the transformer network is the attention head. The attention head evaluates the hidden states in *H* with one another to obtain an output score **z**. The superscript denoting the layer and segment of the following equations are dropped as identical operations are performed at each layer and segment.

The query (**q**), key (**k**) and value (**v**) vectors are calculated from the hidden state **h**:

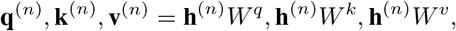

where *W^q^, W^k^*, 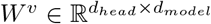 and **q**, **k**, 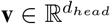. The **q** and **k** vectors are used to obtain a score between two hidden states, expressing their relevance with one another in regard to the information represented by **v**.

For each hidden state at position *n* of the segment, the attention score **z** is calculated by evaluation of its vector **q** with the **k** and **v** vectors derived from the other hidden states in the segment:

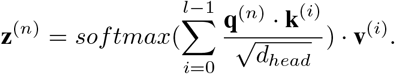

The *softmax* function is used to rescale the weights assigned to the vectors **v** to sum to 1. Division by the square root of *d_head_* is applied to stabilize gradients [35].

The calculation of attention within the attention head is performed in parallel for all hidden states in *H*:

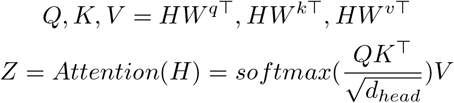

where *Q, K*, 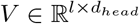 and 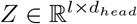. Here, the *softmax* function is applied to every row of *QK*^⊤^.

To increase the capacity of the model, the input is processed by multiple attention heads (*n_head_*) present within each layer, each featuring a unique set of weight matrices *W^q^, W^k^, W^v^* – optimized during training. Having multiple sets of *W^v^*, *W^q^* and *W^k^* allows the model to extract multiple types of information from the hidden states.

The output of the multi-head attention unit is obtained by concatenation of all *Z* matrices along the second dimension and multiplication by *W^m^*. This creates an output with dimensions equal to *H*:

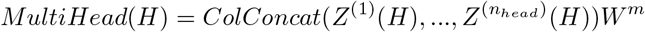

where 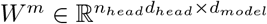.

Next to the information content of the input, positional information of the hidden states is relevant towards the calculation of attention. Unlike the majority of other machine learning methods in the field (e.g. linear regression, convolutional/recurrent neural networks), the architecture of the model does not inherently incorporate the relative positioning of the inputs. Positional information is added by introduction of a bias related to the vector representation and relative distance of the evaluated hidden states [9].

#### 3.1.2 Recurrence

To process the full genome sequence, a recurrence mechanism is applied, as described by Dai et al. [9]. This allows for the processing of a single input (i.e. the genome) in sequential segments of length *l*. In contrast with calculation of the attention heads described in the previous section, only upstream hidden states are used to calculate the output of **h**. In each layer, hidden states [**h**^(*n*+1)^,…, **h**^(*l*−1)^] are masked when processing **z**^(*n*)^, *n* ∈ [0, *l*[.

In order to extend the receptive field of information available past one segment, hidden states of the previous segment *s* − 1 are accessible for the calculation of **h**^(*s,t*+1)^. The segment length *l* denotes the span of hidden states used to calculate attention. Therefore, *H*^(*s,t,n*)^, representing the collection of hidden states used for the calculation of multi-head attention at position *n* in layer *t*+1 of segment *s*, consists of *l* hidden states spanning over segment *s* and *s* − 1:

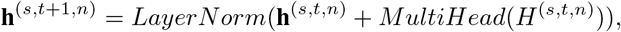

where,

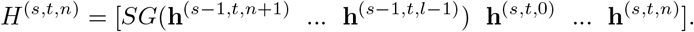

SG denotes the stop-gradient, signifying that during training, no weight updates of the model are performed based on the partial derivatives of given hidden states with the loss. This alleviates training times, as full backpropagation through intermediary values would require the model to retain the hidden states from as many segments as there are layers present in the model, a process that quickly becomes unfeasible for a model with a large segment length or high amount of layers. Figure 1 gives an overview of the model architecture adopting the recurrence mechanism.

**Figure 1:**
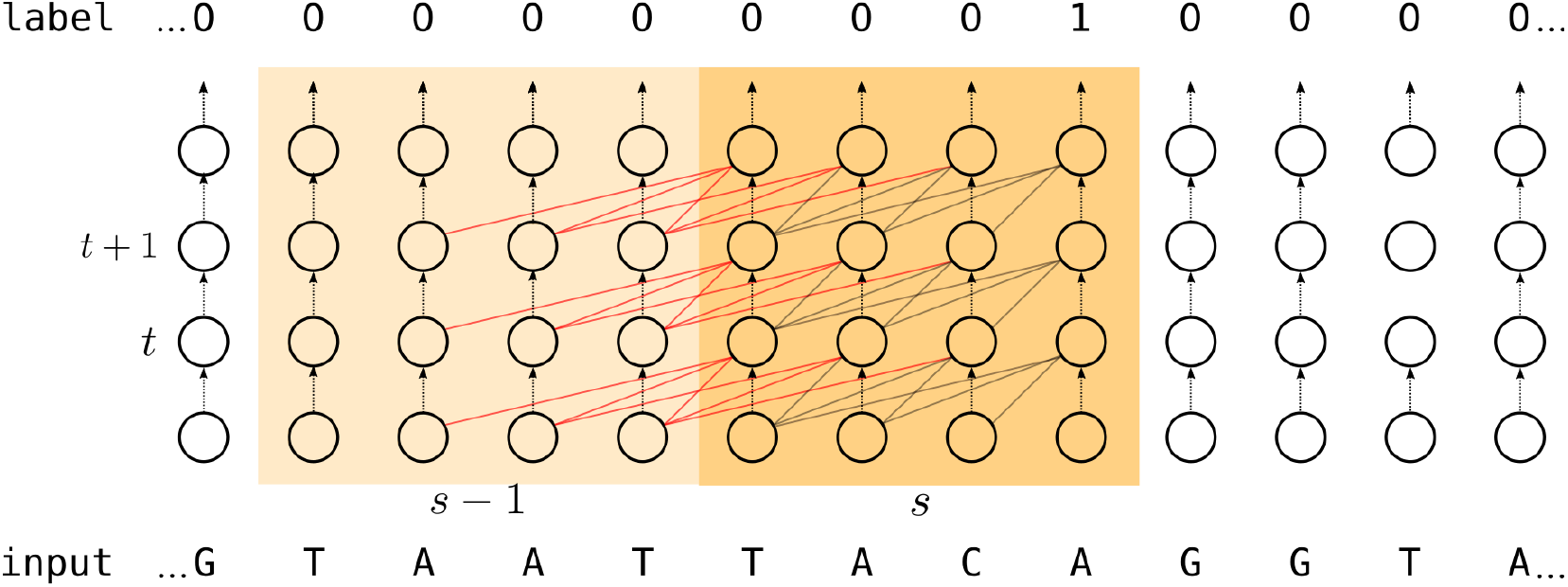
Simplified representation of the architecture of the implemented transformer network. The model processes the genomic strand through segments *s* of length *l* to predict the label. Data is sequentially processed in parallel through *k* layers. Within each segment, outputs are derived from the combination of data from the previous hidden states of the previous layer (grey connections). Cached data from the previous segment are also used, albeit no backpropagation is possible during training (red connections)

### 3.2 Extension: Convolution over *Q, K* and *V*

Important differences exist between the input sequence of the genome and typical natural language processing tasks. The genome constitutes a very long sentence, showing low contextual complexity at input level. Indeed, only four input classes exist. Attention is calculated based on the individual hidden states **h**. For example, in the first layer, hidden states of the segment solely contain information on the nucleotide classes. In previous studies, meaningful sites and regions of interest on the genome are specified by (sets of) motifs from neighboring nucleotides.

To expand the information contained in **q**, **k** and **v** to represent k-mers rather than single nucleotides, a 1D-convolutional layer is implemented that convolves over the **q**, **k** and **v** vectors derived from neighboring hidden states, present as adjoining rows in *Q*, *K* and *V*. The length of the motif, k-mer or kernel is denoted by *d_conv_*.

To ensure that the dimensions of **q**, **k** and **v** remain identical after the convolutional step, as many sets of weight kernels are trained as *d_head_*. Furthermore, through padding, the size of the first dimension of the matrices *Q*, *K* and *V* can be kept constant. Applied on **q** we get:

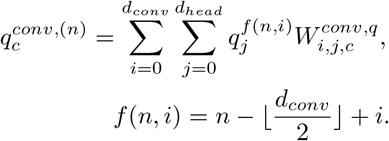

where *c* ∈ [0, *d_head_*[and 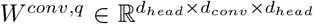. *W^conv,q^* is the tensor of weights used to convolve **q**. Applied on the *Q* matrix the operation is represented as:

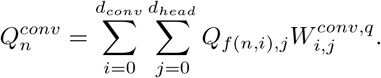

A unique set of weights is optimized to calculate *Q^conv^*, *K^conv^* and *V ^conv^* for each layer. To reduce the total amount of parameter weights of the model, identical weights are used to convolve *Q*, *K* and *V* for all attention heads in the multi-head attention module. Figure 2 gives a visualization of the intermediate results and mathematical steps performed to calculate attention within the attention head of the extended model.

**Figure 2:**
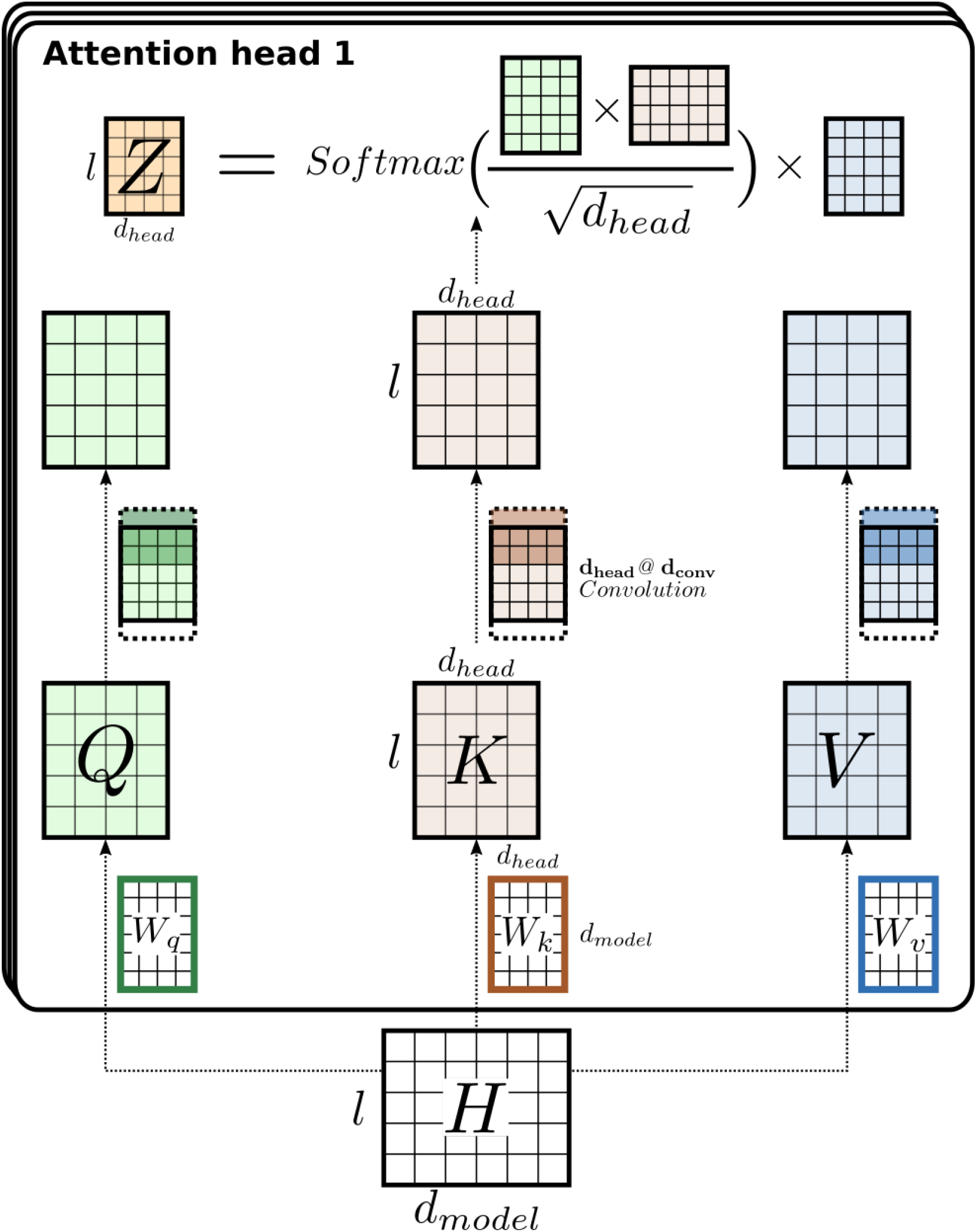
An overview of mathematical operations performed by the attention head to calculate attention **z** for each hidden state **h**. Operations to calculate attention are performed in parallel for *l* hidden states (*H* → *Z*). The query **q**, **k** and **v** vectors are obtained through multiplication of *H* with the model weight matrices *W^q^*, *W^k^* and *W^v^*, resulting in the *Q*, *K* and *V* matrix. A single convolutional layer applied on *Q*, *K* and *V*, using as many kernels as *d_head_*, results in the transformation of the individual **q**, **k** and **v** vector representations of each input to be derived from the **q k** and **v** vectors of *d_conv_* bordering nucleotides. Attention *Z* is thereafter calculated. Through padding, dimensions of *Q*, *K* and *V* are kept constant before and after the convolutional layer. Matrix dimensions are given along the edges of the matrices. The schema is kept simple for better understanding and does not include the relative position encodings added to the *Q* and *K* matrix, nor does it incorporate the recurrence mechanism.

## 4 Experiments and analysis

### 4.1 Experimental setup

To highlight the applicability of the new model architecture for genome annotation tasks, it has been evaluated on multiple prediction problems in *E. coli*. All tasks have been previously studied both with and without deep learning techniques. These are the annotation of Translation Start Sites (TSSs), specifically linked to promoter sites for the transcription factor *σ*^70^, the Translation Initiation Sites (TISs) and N4-methylcytosine sites. The genome was labeled using the RegulonDB [31] database for TSSs, Ensembl [8] for TISs and MethSMRT [40] for the 4mC-methylations. These data sets contain the positively labeled positions on the genome, with all other positions being negative.

For every prediction task, the full genome is labeled at single nucleotide resolution, resulting in a total sample size of several millions. A high imbalance exists between the positive and negative set, the former generally being over four orders of magnitudes smaller than the latter. An overview of the data sets and the resulting sample sizes when labeling the genome are given in Table 1.

**Table 1:**
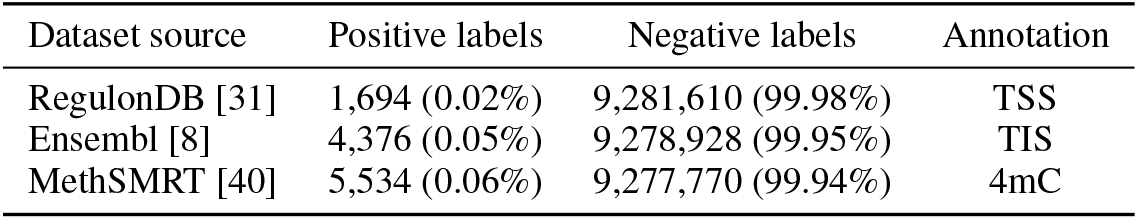
Overview of the data set properties used in this study. From left to right: the name of the database, positive labels, negative labels and annotation task performed (TSS: *σ*^70^ Transcription Start Site; TIS: translation Initiation Site; 4mC: 4mC methylation sites). All data sets are derived from *E. coli* MG1655 (accession: NC_000913.3, size: 9,283,304 nucleotides).

It can be important to include information located downstream of the position of interest to the model, as it is relevant for the prediction of the site. For all three prediction problems, the nucleotide sequence up to 20 nucleotides downstream of the labeled position is regarded as relevant. In order to make this information accessible to predict the label at position *n*, the labels can be shifted downstream. In accordance with the downstream bound taken by recent studies for the annotation of all three annotation tasks [17, 7, 34], labels have been shifted downstream by 20 nucleotides, placing them at position +20 from their respective nucleotide site.

The receptive field of the model is defined as the nucleotide region that is linked indirectly, through calculation of attention in previous layers, to the output. As the span of hidden states used to calculate attention is equal to the segment length *l*, the receptive field has a nucleotide coverage that is equal to the multiplication of the amount of layers with the segment length (*k* × *l*). After shifting the labels downstream, the range of the nucleotide sequence within the receptive field of the model at position *i* is delimited as ]*i* − *k* × *l* + 20, *i* + 20].

As the model sequentially iterates the genome, the training, test and validation set are created by splitting the genome at three positions that constitute 70% (4,131,280–2,738,785), 20% (2,738,785– 3,667,115) and 10% (3,667,115–4,131,280), respectively. An identical split was performed for each of the prediction tasks. Split positions given are those from the *RefSeq* database and, therefore, include both the *sense* and *anti-sense* sequence within given ranges. The model is optimized using the cross-entropy loss and Adam step update algorithm. Surprisingly, weighing of the loss in order to account for the imbalance in class distributions did not have an effect on the performances, and was therefore not done. Model training is stopped when a minimum loss on the validation set is reached. All performance metrics listed are obtained on the test set. Models were trained and evaluated using a single GeForce GTX 1080 Ti and programmed using PyTorch [28].

Hyperparameter tuning was performed using random grid search. To better handle the sheer amount of hyperparameters, models were optimized using a reduced training set (30%) for all three problems. Hyperparameter sets were evaluated based on the minimum loss on the validation set. Due to the limited amount of input and output classes, it quickly became obvious that the required vector size of the hidden states (*d_model_*) was much smaller (32) as compared to the one used for natural language processing tasks (512). Similar observations were made to the overall complexity of the model, reflected by e.g. the amount of layers and attention heads. Given a minimal capacity, the model returned stable performances, with more complex models only increasing processing times. The final hyperparameter set results in a model size that is as minimal as possible, without negatively inhibiting the performances. This set proved to work well on all annotation tasks. As such, only a single model architecture has been used to evaluate all three annotation tasks. Relevant model parameters and optimization parameters are listed in Table 2.

**Table 2:**
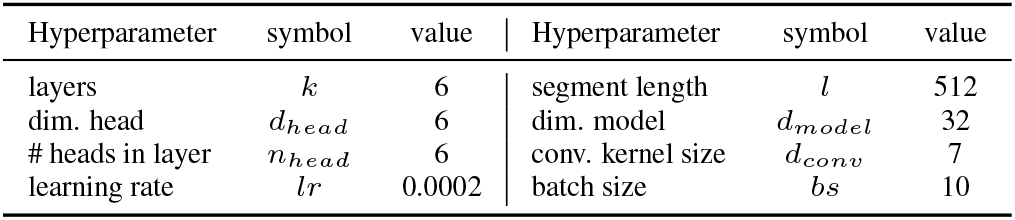
Overview of the hyperparameters that define the model architecture. A single set of hyperparameters was selected to train a single model that showed to work well on all prediction tasks.

The Area Under the Receiver Operating Characteristics Curve (ROC AUC) is the metric selected to evaluate the model performance. This measure is commonly used for binary classification and is independent of the relative sizes of the positive and negative sets.

### 4.2 Improvement by convolution over Q, K and V

The use of nucleotide embeddings as inputs to the transformer network is an important difference with natural language processing, where nucleotides, featuring only four input classes, feature lower contextual complexity than words. Essentially, the equivalent for words are motifs or k-mers present within the DNA sequence. These are of importance towards the biological process of the prediction task due to their affinity towards the domains of related proteins.

In order to investigate ways to improve the model, the reduction of the input and output resolution of the model was first investigated. The use of k-mers as inputs increases the information content of the input embeddings and can facilitate the detection of relevant motifs. The use of k-mers results in a higher amount of input classes (i.e. 4^*k*^) and speeds up processing time as the sequence length is divided by the k-mer size, albeit at the cost of a decreased output resolution of the model predictions.

The reduced performances resulting from applying k-mer inputs underline the disadvantages of this approach. First, different unique sets of k-mers can be used to represent the DNA sequence, determined by the position where splits are performed. Therefore, motifs of relevance to the prediction problem can be represented by multiple sets of input classes. Given the low amount of positive samples of the investigated prediction problems, all possible input class combinations that are of importance are more likely to be only present in either the training, validation or test set. Therefore, higher values of *k* quickly results in the overfitting of the model on the training set.

The high similarity between the k-mers with largely equal sequences (e.g. AAAAAA and AAAAAT) can be mapped through the embedding of the input classes, obtained by Equation 1. Embedding representations for each input class can either be optimized during training or before. In case of the optimization during training, embeddings are in function of the prediction problem (i.e. loss on the labeling), an option less suited for a setting with a small positive set and high amounts of input classes. For the unsupervised setting, vector embedding can be mapped to the input classes using plethora of prokaryotic genomes. This has been done using the word2vec methodology for all classes present in a 3-mer or 6-mer setting, resulting in a slight improvement of the model performances, albeit lower than the performance of the model trained at single-nucleotide resolution [26].

Alternatively, the use of a convolutional layer in the attention heads of the neural network has been investigated. Theoretically, the implementation of a convolutional layer extends the the information embedded in **k**, **q** and **v** to be derived from *d_conv_* neighboring hidden states, without changing the input/output resolution of the model.

This extra step increases the contextual complexity within the attention heads without extending training times substantially, albeit at an increase of the number of model weights. An overview of the mathematical steps performed in the adjusted attention head is shown in Figure 2. To evaluate, performances were compared for different sizes of *d_conv_* for the prediction of TSSs, TISs and methylation sites.

Application of the transformer network with no convolutional layer results in a ROC AUC of 0.919, 0.996 and 0.951 for the annotation of TSSs, TISs and 4mC methylation sites, respectively. Addition of the convolutional layer improves these performances, where *d_conv_* = 7 gives the best results for all three annotation tasks. The increased performance score is most notable for the annotation of TSSs, showing an improvement from 0.919 to 0.977, where the difference with a perfect score is almost divided by four. Performances are given in Table 3. Additionally, the total amount of model parameters and average durations to iterate over one epoch are given.

**Table 3:** Comparison between the performance of the transformer model with different kernel sizes of the convolutional layer on the test sets of the genome annotation. *d_conv_* = 0 constitutes the transformer network with no convolutional step. For all settings, *d_conv_* = 7 results in the best performances. For the annotation of *σ*^70^ Transcription Start Sites (TSSs), the performances for all evaluated *d_conv_* are given, similar to those given in Figure 3. Performances are evaluated using the Area Under the Receiver Operating Characteristics Curve (ROC AUC). For each setting, the total amount of model weights and time (in seconds) to iterate one epoch during training is given.

| Task | $d_{conv}$ | Epoch time (s) | Model weights | ROC AUC |
| --- | --- | --- | --- | --- |
| TSS | 0 | 337.5 | 185,346 | 0.919 |
| TSS | 3 | 364.8 | 241,218 | 0.966 |
| TSS | 7 | 371.0 | 314,946 | 0.972 |
| TSS | 11 | 379.2 | 388,674 | 0.970 |
| TSS | 15 | 383.8 | 462,402 | 0.973 |
| TIS | 0 | 337.5 | 185,346 | 0.996 |
| TIS | 7 | 371.0 | 314,946 | 0.998 |
| 4mC | 0 | 337.5 | 185,346 | 0.951 |
| 4mC | 7 | 371.0 | 314,946 | 0.985 |

The loss curves on the TSS prediction task for all model types are given in Figures 3, and show a stable convergence of the loss for both the training and validation set for *d_conv_* < 7. In contrast, for *d_conv_* > 7, the loss curve on the validation set shows a stronger similarity to a hyperbolic, a pattern that clearly demonstrates overfitting of the model to the training set due to the increased amount of parameter weights in the model.

**Figure 3:**
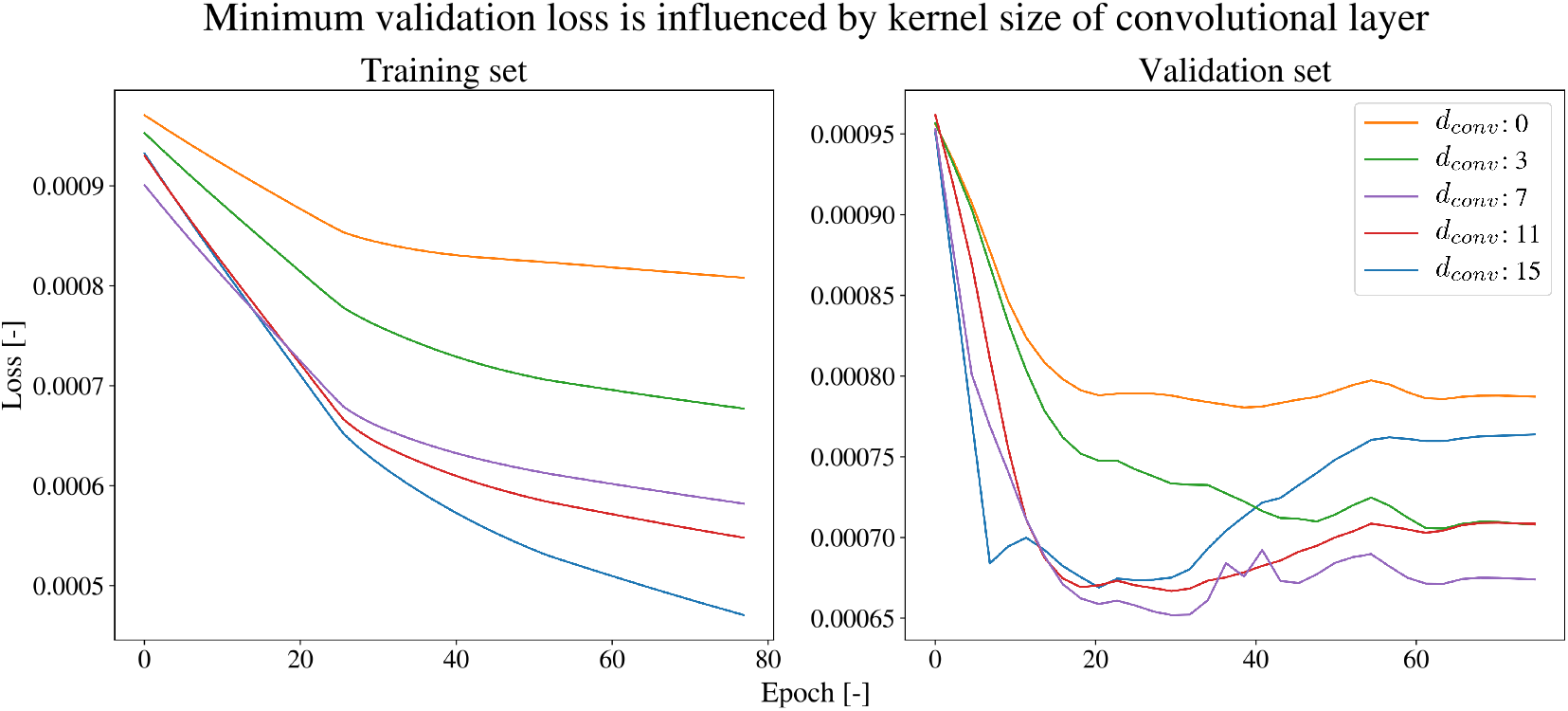
The smoothed loss on the training and validation set of the *σ*^70^ TSS data set for different values of *d_conv_*. In line with the loss curve of the validation set, the best performance on the test set was obtained for *d_conv_* = 7. It can be observed that higher values of *d_conv_* quickly results in overfitting of the model while lower values result in convergence of both the training and validation set at a higher loss.

Importantly, the capacity of the model can be increased through selection of the total number of layers *k*, the amount and dimension of the attention heads *n_head_* and *d_head_* and the dimension of the hidden states *d_model_*. Nonetheless, during hyperparameter tuning, increasing the capacity of the model did not further improve the performance on the test set. The addition of the convolutional layer is thus a necessary enhancement of the model in this setting.

### 4.3 Selection of *l_mem_*

The parameter *l_mem_* denotes the amount of last-most hidden states of the previous segment (*s* − 1) stored in memory, thereby delimiting the amount of hidden states made accessible for the calculation of attention in layer *s*. Traditionally, *l_mem_* is set to *l* during training time, ensuring the calculation of attention at each position in segment *s* to have access to *l* hidden states (as stored in *H*^(*n,t,s*)^) [9]. As *l_mem_* determines the shapes of *H*, *K* and *V*, it is a major factor influencing the memory requirement and processing time of the model.

In order to reduce training times to enable the applicability of the framework for larger genomes, data from several models (*d_conv_* = 7) for the annotation of *σ*^70^ TSSs was collected for different values of *l_mem_* during training (denoted by 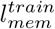). Additionally, as the segment length *l* is not tied to the model weights and can be altered after training, performance metrics for different segment lengths of the model for the annotation of the test set (*l^test^*) were also obtained. 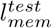 is always set equal to *l^test^*.

The processing time to iterate the genome is reduced by halve for 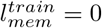. Figure 4 shows the loss on the training and validation set in function of time for the different values of 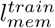. Interestingly, after training of 75 epochs, lower losses on the training set are obtained for lower values of 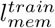. No backpropagation is performed through the hidden states of the previous segment (see Section 3.1.2), even though the above elements contribute to the loss during training. The inability to properly update the weights of the model in function of hidden states from previous segments are a likely cause for the slower convergence of the training loss for 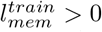. Therefore, processing times are reduced both by the reduction of epoch time and the fewer epochs required until a minimum on the validation loss is obtained.

**Figure 4:**
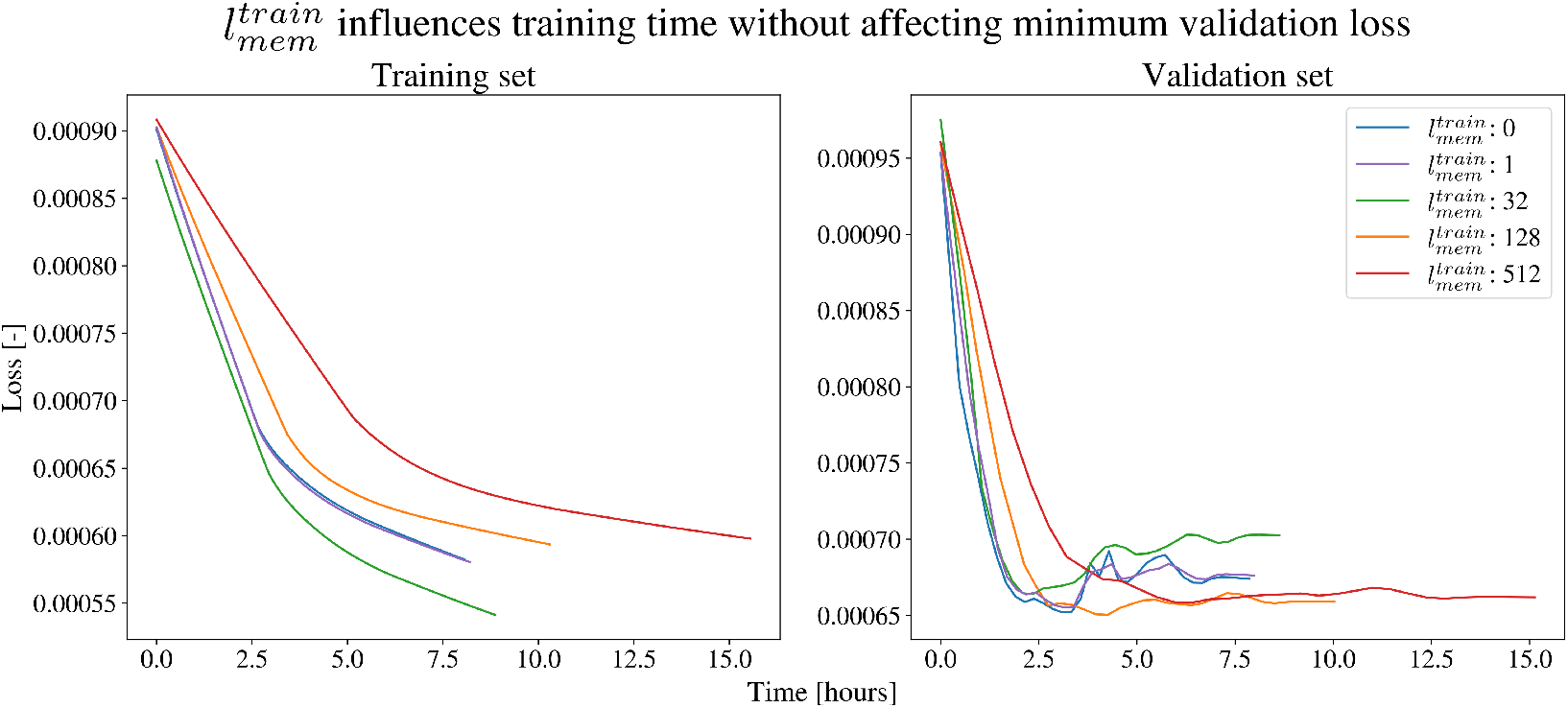
The smoothed loss on the training and validation set of the *σ*^70^ TSS data set for different values of 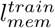. The losses are given w.r.t. training times. All settings were trained for 75 epochs. While increased values of *l_mem_* strongly influence the convergence time of the loss for both the training and validation set, it does not alter the minimum loss on the validation set.

Table 4 shows the ROC AUC performances and training times for the annotation of *σ*^70^ TSSs. Models trained for 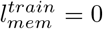 are not penalized in their performance on the test set. In contrast, for all values of 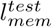, higher performances as compared to the traditional setting 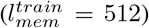 are obtained. The strong variation of hidden states applied to calculate attention for **h**^*n*^, ranging between 1 to 512 and dependent on the position of *n* within *s*, does not negatively influence the performance. The discussed variation might in fact contribute to regularization of the model weights, given the more stable results of the model on varying values of *l^test^*.

**Table 4:**
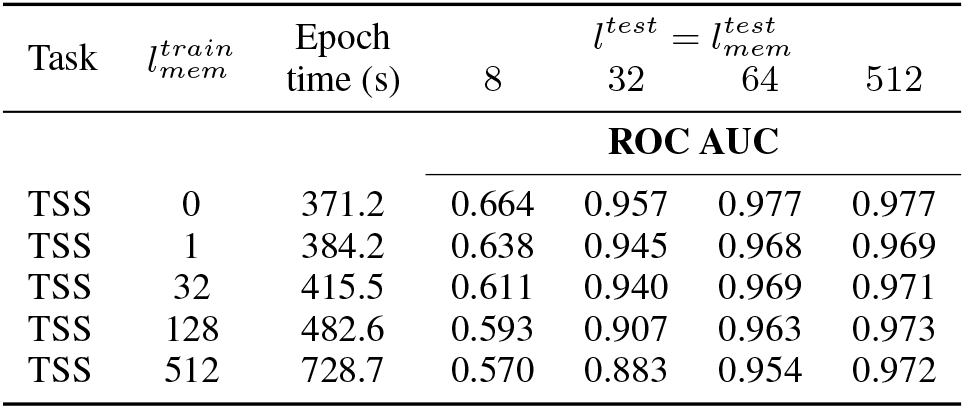
Performances, given as Area Under the Receiver Operating Characteristics Curve (ROC AUC), of the transformer model on the test set for the prediction of *σ*^70^ Transcription Start Sites (TSS) using different values of 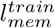 and *l^test^*. For each setting, *l_train_* = 512 and 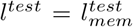. The time (in seconds) to iterate one epoch during training is also given.

The receptive field of the models predictions for 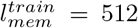 spans 3,072 nucleotides (*l* × *k*), a region multiple times larger than the circa 80 nucleotides window used in previous studies [34][23][29]. Reduction of *l^test^* can offer insights into the DNA region relevant towards the prediction task. This is illustrated by the performances of the model for *l^test^* equal to 64 and 512, where the reduction of the receptive field to 384 nucleotides does not negatively influence performances (for 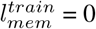), revealing the excluded region to be of no importance towards the identification of a TSS. Overall, given the influence of the segment lengths on both the training time and performance, a closer look into the behavior of the model for varying values of *l* and *l_mem_* should be made for the genome annotation tasks. In this study, the model parameters *l^train^* = 512, 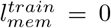 and 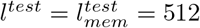 proved to work best for all three annotation tasks.

### 4.4 Benchmarking

As a final step, the proposed transformer model (*d_conv_* = 7, 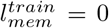) has been evaluated with the studies reporting state-of-the-art performances on all three annotation tasks. For each setting, the same annotations have been applied to train and evaluate all models, meaning that the positive set between these studies is identical. Differences exist between the negative sets, as our method processes the full genome. This is in contrast to the use of a subsampled negative set by recent studies [29, 24, 36, 23, 34, 6, 29]. In the majority of studies, custom sampling methods are used, where the samples of the negative set is not publicly available[23, 29, 34, 17, 6]. As the transformer architecture iterates the genome sequentially, the creation of a training, test and validation set have are obtained by slicing the genome at three points. This results in neighboring nucleotides being grouped in the same set, which might introduce a bias. However, it was made sure that the relative frequencies of the input and output classes were identical for all sets. No bias was detected, as the performances of the transformer model proved to be robust after training and evaluating the model using various sets.

Uniquely, the transformer model processes the full negative set for training and evaluation purposes. Several metrics used to represent the performance are directly influenced by the (relative) sizes of the negative and positive sets. These include the accuracy and Area Under the Precision-Recall Curve (PR AUC). Furthermore, the sensitivity (recall), specificity, precision and Matthew Correlation Coefficient are all metrics that depend on the threshold used to group the model output probabilities into positive and negative predictions. This threshold can be selected to maximize any metric, such as the accuracy, recall or precision.

Figure 5 shows the effect of subsampling the negative set on the performance metrics and the selection of the optimal threshold to delineate the positive from the negative predictions. In the first setting, 1500 instances are sampled from the positive and negative output distributions, representing the models ability to categorize both classes. The ROC AUC and PR AUC are both 0.983. The optimal threshold returns a precision and recall of 0.94 and 0.93, respectively. Evaluating the performance of the model for an increased negative set size with 9,000,000 instances, sampled from the same distribution, reveals two problems: the PR AUC performance measure is decreased to 0.25 as a direct result of false positives, and the use of the threshold selected from the first set-up does not properly balance the trade-off between false negatives and false positives. Specifically, the accuracy, precision and recall equal to 0.933, 0.0046 and 0.93, as compared to 0.999, 0.713 and 0.136 for a threshold optimized for the accuracy on the second setting.

**Figure 5:**
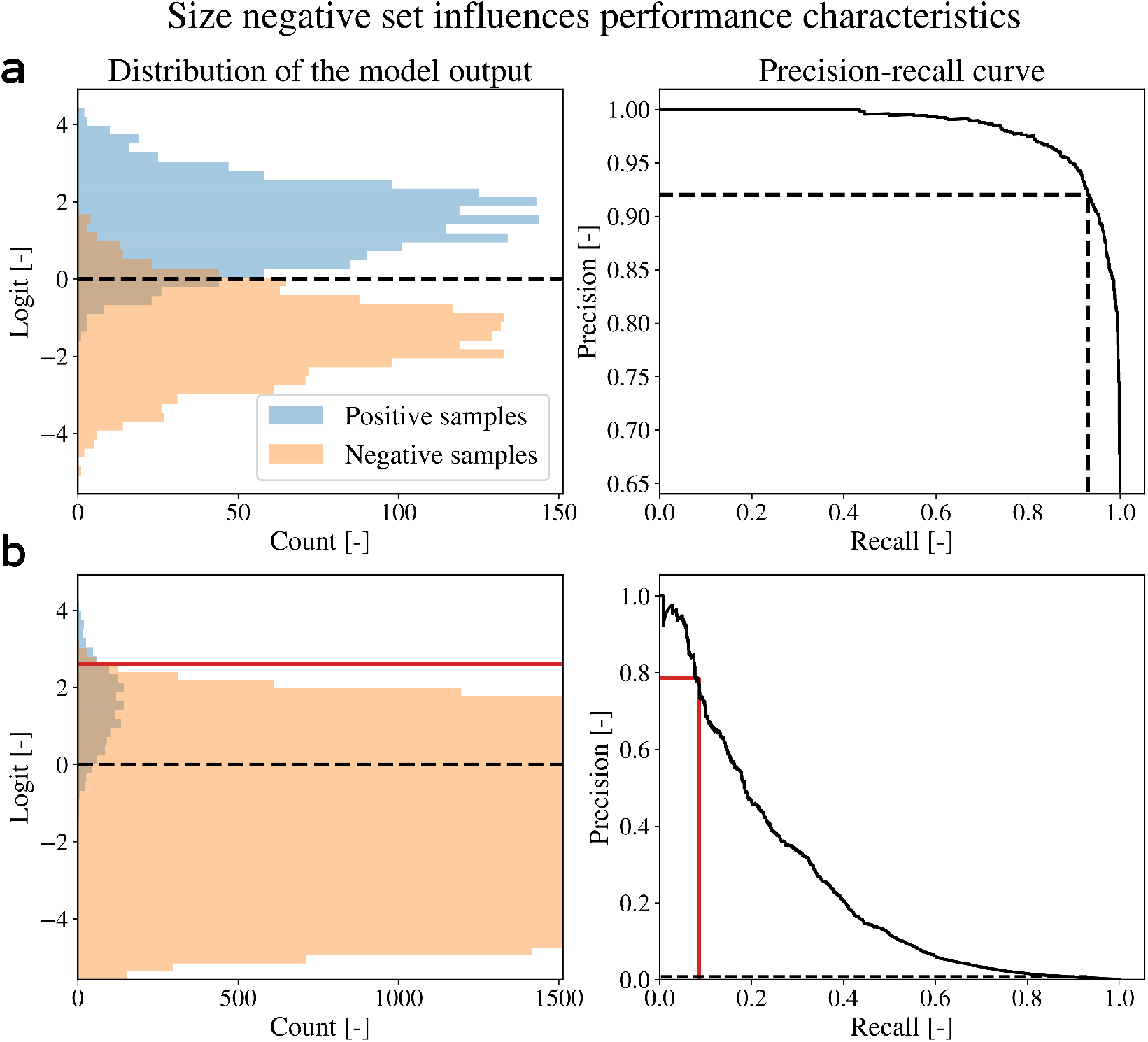
Illustration of varying model performance characteristics as an effect of different sample sizes. The distributions of the outputs and precision recall-curves are given. (**a**) An equal positive and negative sample size of 1500 gives an optimal accuracy of 0.936. The coinciding threshold selected to optimize accuracy (gray striped line) gives a precision of 0.94 and recall of 0.93. (**b**) Given the same distribution characteristics but with negative sample size of 9,000,000, performance of the PR AUC is drastically reduced (0.25) as a result of false positives. Moreover, the threshold selected to optimize accuracy is adjusted (green full line). The resulting accuracy, precision and recall for the new setting are 0.999, 0.711 and 0.136. Using of the threshold from the previous setting gives the scores of 0.933, 0.0046 and 0.93.

The ROC AUC metric is independent to the relative sizes of the positive and negative set. Scores in Table 5 list the ROC AUC scores of recent studies claiming state-of-the-art performances. For CNNs, the model architectures have been implemented as described in each study to attain the ROC AUC performance metrics. As such, performances have been obtained from these models using exactly the same positive, negative, training, validation and test set used to obtain performances on the transformer model.

**Table 5:**
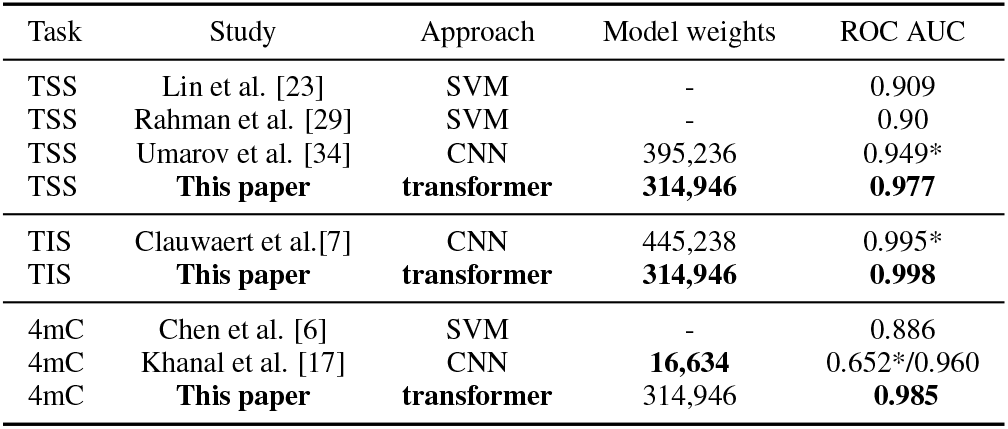
Performances, given as Area Under the Receiver Operating Characteristics Curve (ROC AUC), of recent studies and the transformer based model on the annotation of *σ*^70^ Transcription Start Sites (TSS), Translation Initiation Sites (TIS) and 4mC methylation (4mC). Performances listed are those reported by the paper, and generally constitute a smaller negative set. Additionally, performances with an asterisk (*) are obtained by implementation of the model architecture and training on the full genome. Applied machine learning approaches include Convolutional Neural Networks (CNN) and Support Vector Machines (SVM). When applicable, the amount of model weights is given.

| Task | Study | Approach | Model weights | ROC AUC |
| --- | --- | --- | --- | --- |
| TSS | Lin et al. [23] | SVM | - | 0.909 |
| TSS | Rahman et al. [29] | SVM | - | 0.90 |
| TSS | Umarov et al. [34] | CNN | 395,236 | 0.949* |
| TSS | **This paper** | **transformer** | **314,946** | **0.977** |
| TIS | Clauwaert et al.[7] | CNN | 445,238 | 0.995* |
| TIS | **This paper** | **transformer** | **314,946** | **0.998** |
| 4mC | Chen et al. [6] | SVM | - | 0.886 |
| 4mC | Khanal et al. [17] | CNN | **16,634** | 0.652*/0.960 |
| 4mC | **This paper** | **transformer** | 314,946 | **0.985** |

The ROC AUC of 0.977, 0.998 and 0.985 for the annotation of *σ*^70^ TSSs, TISs and 4mC methylation sites represent a substantial improvement of the performances obtained by previous studies. In essence, the improvement of the ROC AUC is substantial as it more than halves the area above the curve (0.949 → 0.977, 0.995 → 0.998 and 0.960 → 0.985). The score for the model implemented in accordance to the work of Khanal et al [17] shows a strong variation with the reported score (0.652/0.960). It was not possible to pinpoint the cause for this discrepancy, but it could be related to the increased size of the negative set. As the transformer model outperforms either, this was not investigated further.

With the exception of the CNN model for 4mC methylation, the amount of model weights is in line with previous neural networks developed. A single architecture for the transformer model (*d_conv_* = 7, 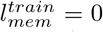) was trained to perform all three annotation tasks, proving the robustness of the novel framework for genome annotation tasks. Implementation of the convolutional layer within the attention heads proved to be required to achieve state-of-the-art results using the proposed attention networks.

## 5 Conclusions and Future Work

In this paper, we introduced a novel framework for full genome annotation tasks by applying the transformer-XL network architecture. To extend the calculation of attention beyond hidden states derived from the nucleotides inputs, of which only four input classes exist, a convolutional layer over *Q*, *K* and *V* was added. As an effect, calculation of relevance (*QK*^⊤^) and linear combination with *V* processes information to be derived from multiple neighboring hidden states. An improvement in predictive performance was obtained, which indicates that the technique enhances the detection of nucleotide motifs that are relevant to the prediction task.

The efficacy of the transformer network was demonstrated on three different tasks in *E. coli*: the annotation of transcription start sites, transcription initiation sites and 4mC methylation sites. In recent studies applying machine learning techniques for genome annotation tasks, the lack of an existing benchmark data set and the existence of custom negative sets hinders the straightforward and clear comparison of existing methodologies. In a balanced setting, the sampled negative set constitutes only a fraction of the negative samples within the genome (e.g. 0.02%–0.1% for TSSs). Therefore, sampling of the negative set makes the trained model susceptible to overfitting due to bad generalization towards the true negative set. Furthermore, application of the full genome for evaluation purposes ensures the resulting performances to correctly reflect the model’s capability in a practical setting. Both the application of the full negative and postive set and the easy construction of the training, test and validation set facilitate future benchmarking efforts.

Models were trained within 2–3 hours. A single iteration over the prokaryotic genome on a single GeForce GTX 1080 Ti takes ca. six minutes. The transformer architecture does not assert the relative positions of the input nucleotides w.r.t. the output label, a property that makes the methodology well-suited for the processing of genome sequences. First, to annotate each position on the genome, inputs only have to be processed once, as intermediary values are shared between multiple outputs. Second, increasing the receptive field of the model, defined through *l* and *l_mem_*, does not require training a new neural network, and is unrelated to the total amount of model parameters. These advantages improve the scalability of this technique. Specifically, a model with a receptive field spanning 3,072 nucleotides (*l* = 512, 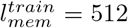) can process the full genome in ca. 12 minutes, as shown in Table 4. Moreover, as shown in this paper, evaluation of the test set for different values of *l*, reveals the minimal receptive field required by the model to obtain optimal performances.

Next to state-of-the-art performances, the application of the transformer-XL architecture and the evaluation on the full genome sequence offer new opportunities. For example, the probability profile of the model along the genome sequence could result in a better understanding of the model and the biological process. In natural language processing, evaluation of attention (*QK*^⊤^) has connected semantically relevant words [35]. Investigation into the profile of the attention scores might pinpoint biological sites of relevancerite in a similar fashion.

Given the success of the models in this study, transformer based networks might show to be valuable in other branches featuring sequence labeling tasks, such as secondary structure prediction of proteins. Due to the size of the eukaryotic genome, application of the technique on these genomes is not feasible at this point. Nevertheless, transformer-based models have not been studied before in this setting, and several areas have potential for further optimization of the training process time. These include the general architecture of the model, batch size, *l^train^*, 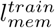, learning rate schedules, etc.

## Acknowledgments

The authors acknowledge the support of Ghent University.

## Code

The code used to train the models is hosted at https://github.com/jdcla/DNA-transformer

